# Succinct Dynamic de Bruijn Graphs

**DOI:** 10.1101/2020.04.01.018481

**Authors:** Bahar Alipanahi, Alan Kuhnle, Simon J. Puglisi, Leena Salmela, Christina Boucher

## Abstract

**Motivation:** The de Bruijn graph is one of the fundamental data structures for analysis of high throughput sequencing data. In order to be applicable to population-scale studies, it is essential to build and store the graph in a space- and time-efficient manner. In addition, due to the ever-changing nature of population studies, it has become essential to update the graph after construction e.g. add and remove nodes and edges. Although there has been substantial effort on making the construction and storage of the graph efficient, there is a limited amount of work in building the graph in an efficient and mutable manner. Hence, most space efficient data structures require complete reconstruction of the graph in order to add or remove edges or nodes.

**Results:** In this paper we present DynamicBOSS, a succinct representation of the de Bruijn graph that allows for an unlimited number of additions and deletions of nodes and edges. We compare our method with other competing methods and demonstrate that DynamicBOSS is the only method that supports both addition and deletion and is applicable to very large samples (e.g. greater than 15 billion *k*-mers). Competing dynamic methods e.g., FDBG (Crawford et al., 2018) cannot be constructed on large scale datasets, or cannot support both addition and deletion e.g., BiFrost (Holley and Melsted, 2019).

**Availability:** DynamicBOSS is publicly available at https://github.com/baharpan/dynboss.

## 1 Introduction

The de Bruijn graph was introduced for assembling genomes from short sequence reads by Pevzner et al. (2001) and since then has been widely-adopted for this and other applications. Given a set of sequence reads of a given individual and an integer *k*, we construct the de Bruijn graph by creating a directed edge for each possible *k*-length subsequence (*k*-mer), labeling the nodes of each directed edge with the longest proper prefix and suffix of the respective *k*-mer, and lastly, gluing together all nodes having the same label. This is the well-known constructive definition of the de Bruijn graph. After construction of the graph, contiguous sequences (contigs) are created by traversing the graph and outputting the sequence(s) corresponding to the traversal. Although many de Bruijn graph representations have been developed – including, Simpson et al. (2009); Conway and Bromage (2011); Chikhi and Rizk (2013); Bowe et al. (2012) – only a few of them are dynamic, meaning that they allow for the addition or deletion of *k*-mers. Crawford et al. (2018), were the first to present a compressed and fully dynamic data structure for storing the de Bruijn graph. Yet, their method, referred to as FDBG, cannot be constructed for large datasets. Hence, existing representations are never both highly-compressed and fully dynamic (e.g. allow for additions and deletions). This is unsurprising since it is generally challenging to store data in a manner that is both space-efficient and mutable.

In 2012, while the development of space efficient representations of de Bruijn graphs was well under way, the so-called *colored de Bruijn graph* was defined. It is described as an intuitive way to model, identify and store the genetic variants of individuals of a given species. One can extend the previous constructive definition of a de Bruijn graph to describe the colored de Bruijn graph. Given *n* sets of sequence reads, *R*_1_, .., *R*_*n*_, we construct the colored de Bruijn graph by labelling each edge with one or more colors, indicating in which set(s) of reads the *k*-mer occurs, i.e., an edge corresponding to *k*-mer *s*_*k*_ contains color *c*_*i*_ if *s*_*k*_ is in *R*_*i*_. Since its introduction, the use of the colored de Bruijn graph was expanded, and they were constructed on increasingly larger, more heterogenous datasets. Cortex, the original colored de Bruijn graph implementation due to Iqbal et al. (2012), was space- and time-inefficient, which limited its use to relatively small, heterogenous datasets.

In order to use colored de Bruijn graphs for finding and storing genetic variants in more complex datasets, a more compact representation was introduced by Muggli et al. (2017). This representation reduces space by building a succinct representation of the “union de Bruijn graph”, which is the de Bruijn graph constructed on all sets of sequence reads (i.e., *R*_1_, .., *R*_*n*_), and then stores the color information separately, as a *n* by *m* matrix of bits, where *m* is the number of *k*-mers. This *color matrix*, typically denoted as *C*, is defined as *C*[*i, j*] = 1 if *k*-mer *j* is contained in *R*_*i*_ and *C*[*i, j*] = 0 otherwise. Hence, this allows the matrix to be compressed and stored independently of the graph. Also several other methods have been developed to further compress, store, and manipulate the color matrix, including Rainbowfish (Almodaresi et al., 2017), Mantis (Pandey et al., 2018), Bloom Filter Trie (BFT) (Holley et al., 2016), and Bifrost (Holley and Melsted, 2019).

The analysis of sequencing data from multiple samples using a colored de Bruijn graph added further pressure to develop dynamic data structures. When a new sample arrives, it needs to be added to an existing colored de Bruijn graph. Furthermore, in order to analyze a large biological dataset in a systematic fashion, it is common to restrict interest to a portion of the samples rather than the complete dataset – which would correspond to removing some samples from a colored de Bruijn graph. For example, several of the most recent papers including (Muggli et al., 2019; Holley and Melsted, 2019) on the colored de Bruijn graph analyze GenomeTrakr data, which are large datasets used for surveillance of antimicrobial resistance pathogens. These datasets contain meta-data describing each sample (dates, sample type, farming vs. clinical data). After constructing the graph on the these data one important type of analysis would be to find all pathogens within a geographical area and/or time period. In order to accomplish this, samples (selected using the meta-data) would have to be deleted from the graph.

The current approaches for building and storing the de Bruijn graph do not have this capability as they are not highly-compressed and dynamic. To address these needs, several methods (Mustafa et al., 2017, 2019; Karasikov et al., 2019) were developed to store the color matrix (or graph annotation) in a manner that is both compressed but also dynamic, meaning it allows for rows and columns of the matrix to be added or deleted. Yet, they do not tackle the problem of implementing a space-efficient, dynamic de Bruijn graph. Hence, Mustafa et al. (2017) state that the construction of a dynamic compressed color matrix is a “step towards implementing a dynamic data structure for indexing large annotated sequence data sets that supports fast query and update operations.” At the time of writing, no established standard tool has filled this niche. BFT (Holley et al., 2016), and Bifrost (Holley and Melsted, 2019) are two bloom filter-based representations of succinct colored de Bruijn graph which support addition of *k*-mers but are not fully dynamic since they cannot support deletion.

### Our contributions

In this paper, we describe a space-efficient dynamic data structure for building the de Bruijn graph in a manner that is practical for large samples and allows for both addition and deletion of *k*-mers. We refer to our method as DynamicBOSS. We accomplish this by bringing dynamism to the (previously static) Burrows Wheeler transform (BWT) based de Bruijn graph representation Bowe et al. (2012) by extending the recent work by Prezza (2017) on dynamic succinct bit vector representations.

We compare DynamicBOSS with all other dynamic de Bruijn graph methods namely, FDBG (Crawford et al., 2018), BFT (Holley et al., 2016), and Bifrost (Holley and Melsted, 2019) as well as the static BOSS data structure using *E*.*coli* and metagenome data. Although FDBG was more efficient at performing addition and deletion, the size of the graph was more than two times the size of DynamicBOSS and it took twice as long to construct. Further, FDBG failed to build the graph on samples with 55 million or more reads. The de Bruijn graph constructed by BFT was more than six times the size of DynamicBOSS, and BFT no longer supports addition (Holley, 2019). The most-recent method, Bifrost required between 3 and 11 times more time to construct the de Bruijn graph, and between 10 and 16 times more memory and up to 6 times more space to store the graph. It required slightly less space (less than 1 bit / *k*-mer) to store the graph on small samples but the space increased significantly with the data size, resulting in DynamicBOSS requiring 2 to 6 times less space on large samples. Moreover, Bifrost did have faster addition time but it does not support deletion and the addition time scaled non-linearly. In conclusion, we show that DynamicBOSS is the only full-dynamic, space-efficient de Bruijn method that can be constructed on large samples.

## 2 Preliminaries

In this section, we go over some of the basic terminology and definitions that will be used throughout this paper.

### Suffix arrays

The suffix array (Manber and Myers, 1993) SA_X_ refers to a succinct representation of the lexicographic ordering of all suffixes of a string X. More specifically SA_X_ is an array SA[1 … *n*] which contains a permutation of the integers [1 … *n*] such that X[SA[1] … *n*] ≺ X[SA[2] … *n*] ≺ … ≺ X[SA[*n*] … *n*]. Here, ≺ denotes lexicographic precedence. Hence, SA[*j*] = *i* if and only if X[*i* … *n*] is the *j*^th^ suffix of X in lexicographical order. The suffix array helps in quickly locating all occurrences of a substring P within X. In order to accomplish this we need to find all suffixes that start with P. Due to the lexicographical order in SA, all such suffixes occur in an interval in SA_X_ which is called suffix array interval and is associated with a pair of integers [*s, e*] denoting the first and last index in SA_X_, corresponding to the suffixes having P as a prefix. Using X and SA_*X*_, *s* and *e* are efficiently found with binary search for P.

### Burrows Wheeler Transform (BWT) and FM-index

BWT (Burrows and Wheeler, 1994) is a reversible permutation of the characters of a string, used originally for compression. Here we give a constructive description of BWT. Given a string X, we first add a $ to the end of X to keep track of end of string. Now similar to the suffix array, if we sort all rotations of X lexicographically, the last column of this sorted list is BWT of X, which we denote by BWT_X_. More specifically, BWT_X_[*i*] = X[SA_X_[*i*] − 1] when SA_X_[*i*] is greater than one and BWT_X_[*i*] = $ when SA_X_[*i*] is equal to one. This simply means, BWT_X_ is the array of characters just to the left of the suffixes in the suffix array SA_X_. Ferragina and Manzini (2005) introduced the FM-index and showed that BWT also can be used for indexing. FM-index is an extension of BWT_X_ with two data structures: (i) array C_X_ such that C_X_[*a*] stores the number of symbols in X that are lexicographically lower than the symbol *a*, and (ii) a ranking data structure Occ_X_ on BWT_X_ such that Occ_X_(*a, i*) keeps the number of occurrences of the symbol *a* in BWT_X_[1, *i*]. Now knowing that the suffix array interval of a string *S* is [*s, e*], we can calculate the interval [*s*′, *e*′] for the string *aS* as follows: *s*′ = C_X_[*a*] + Occ_X_(*a, s* − 1) and *e*′ = C_X_[*a*] + Occ_X_(*a, e*) − 1. In order to compute the occurrences of a substring P within the string X, above calculation is iteratively performed |P| times and since each operation requires constant time, *s* and *e* – first and last indexes of the interval containing P–are found in *O*(|P|), independent of the size of X.

## 3 Dynamic BOSS

In this section, we introduce DynamicBOSS which is a compressed data structure for the de Bruijn graph that supports the addition and deletion of edges and nodes. We begin by describing how to construct and store the de Bruijn graph using BWT and then show how it can be extended to support dynamic operations.

### 3.1 Static BOSS

Given a de Bruijn graph *G* = (*V, E*), we refer to the label of an edge *e* ∈ *E* as the *k*-mer corresponding to it, and denote it as elabel(*e*). Similarly, we refer to the label of a node *v* ∈ *V* as the (*k* − 1)-mer corresponding to it, and denote it as nlabel(*v*). First, we notice that one method to store *G* is to simply store all edges (*k*-mers) in an array sorted in co-lexicographic (colex) order by the nlabel of their ending nodes, where ties are broken by their starting nodes. In order to traverse *G* from a given node, we can find all incident nodes by looking at the *k*-th character and their position in the array via binary search. Although, this simple representation fully specifies *G* and allows for insertions and deletions of nodes and edges, it requires *O*(*k*|*E*|)-space, which is prohibitively large. Next, we show how to represent *G* using a significantly less space using BWT but also allow for insertions and deletions of nodes and edges. We will use this simple representation as a stepping stone for our more complicated representation.

Hence, we begin by showing how to transform this simple representation into one that uses far less space, i.e., one bit vector, one character array, and one integer array. To accomplish this we use BWT. We first pad the input so that each node has at least one incoming edge and one outgoing edge. Hence each node with no incoming edge is shifted and prepended with $ signs to create the required incoming dummy edges (*k*−1 each)–recall that *k* − 1 preceding incoming edges are needed to retrieve the label of each node. For instance in Figure 1, the dummy edges $$$C, $$CG and $CGT are added to make the *k* − 1 = 3 preceding incoming edges of the node CGT which did not have a real incoming edges. Note that after adding the required incoming dummy nodes and edges, the node $$…$ is always the first node in the colex oreder and exceptionally needs no incoming edges. Next, we add one succeeding dummy edge for each node that does not have an outgoing edge by appending a $ to its label. For instance, in Figure 1 the node GAT had no real outgoing edges hence we added GAT$.

**Fig. 1:**
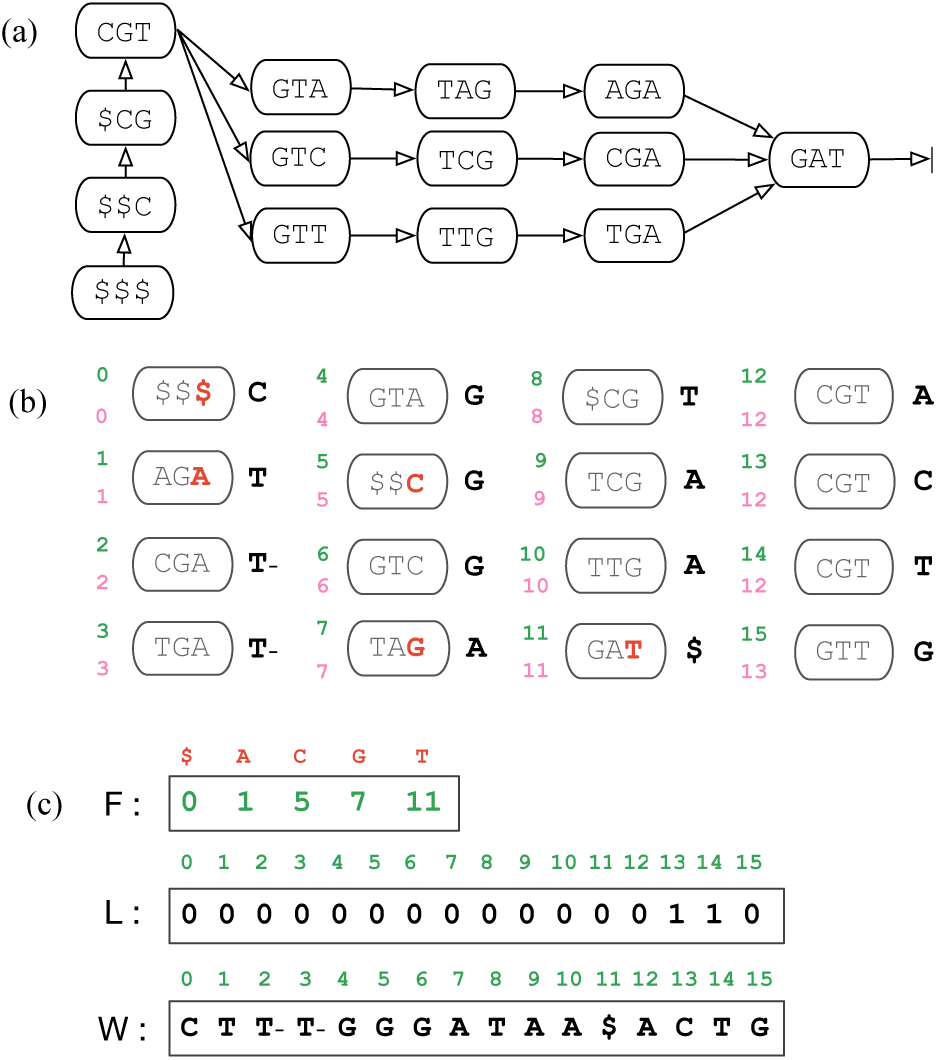
(a) The de Bruijn graph for sequences CGTAGAT, CGTCGAT, CGTTGAT. Note that the dummy edges are $$$C, $$CG, $CGT and GAT$. (b) All edges of the de Bruijn graph are sorted in colex order of the node labels (nlabel). The node index and the edge index are shown in pink and green respectively on the left side and the edge label is shown on the right side of each edge. The first occurrence of each node label is shown in red. (c) The BWT-representation of the de Bruijn graph. Vector F is an array of length five that stores the position (edge index) of the first occurrence of each node label. Bit vector L stores 0 at the position of first outgoing edge of each node and 1 at the position of every other outgoing edge. W is the last column of the sorted elabel’s with the addition that we add a minus sign to the character if it is not the first incoming edge to its target node. For instance we add minus to edges CGAT and TGAT since they are the second and third incoming edges to the target node GAT.

We then sort the list of all elabel’s (dummy edges included) in colexorder by nlabel with all duplicate entries removed, i.e., the list of elabel’s is now sorted by the first *k* − 1 characters. We refer to this ordering of the nodes/edges as *BOSS-sorted*.

Notice that the last column of the nlabel’s (second last in the elabel’s) is sorted in alphabetical order. We refer to this vector as F; thus, F[*i*] is the last character of the node label of node *i*. The vector F is easily compressed by employing an array of length five that stores the position of the first occurrence of each character in that column. Next, with the goal of not storing the first *k* − 2 characters of the elabel’s (since we only need the two last columns of elabels) we add some auxiliary information which enable us to support rank and select. We add a bit vector L which stores 0 at the position of the first outgoing edge of each node and 1 at the position of every other outgoing edge. Next, we define a vector W as the last column of the sorted elabel’s with the addition that we add a minus sign to the character if it is not the first incoming edge to its target node; otherwise, it remains unchanged. For instance in Figure 1 we add minus to edges CGAT and TGAT since they are the second and third incoming edges to the target node GAT. Hence, W is defined on the alphabet {$, *A, C, G, T, A*-, *C*-, *G*-, *T* -}.

Therefore, L, W, and F is an equivalent representation to the previous (simple) one but uses far less space since the first *k* − 2 characters of the elabel are eliminated. This representation, which we refer to as BOSS, is due to Bowe et al. (2012). Figure 1 illustrates this representation. We note that we refer the *node label* of a node as the last character of the nlabel, which is indirectly stored in F. Similarly, the *edge label* of an edge is defined as the last character of elabel, which is stored in W. Also, we refer to the *edge index* of an edge *e* as the position of its elabel in BOSS-sorted order. Similarly, the *node index* of a node *v* is the position of its nlabel in BOSS-sorted order. This is shown in Figure 1(b).

### 3.2 From Static to Dynamic

Recently, many dynamic, compressed bit vectors have been proposed (Mäkinen and Navarro, 2006; Grossi et al., 2013; Navarro and Nekrich, 2014; Klitzke and Nicholson, 2016; Cordova and Navarro, 2016; Álvarez García et al., 2019). In our work, we replace L and W with the compressed, dynamic bit vector implementations of Prezza (2017). With these replacements, there exists an increase in the space and time complexity to store and access the vectors. In particular, the time to access an element of W and L is O(log |Σ| log *n*) and O(log *n*), respectively, where *n* is the number of elements and |Σ| is the alphabet size.

### 3.3 Graph Operations

Below is a partial list of functions that our method and the original BOSS data structure support. All functions use rank, select, and access on W and L and access on F. We refer the reader to Bowe et al. (2012) for detailed discussion of these functions. The complexity of all of the operations on W is O(log |Σ| log |*E*|) and on L is *O*(log |*E*|), where |*E*| is the number of edges and |Σ| is the alphabet size (Prezza, 2017), which is equal to 5 in DynamicBOSS. Lastly, we note the access time of array F is constant.

- index(*e*) returns the index *i* of the edge whose label is equal to *e*. The value −1 is returned if this edge is not in the graph.
- index-node(*u*) returns the index *i* of the first edge of the node whose label is equal to *u*. The value −1 is returned if this node is not in the graph.
- edge(*u*) returns the index of the first outgoing edge of node *u*.
- node(*e*) returns the index of the source node of edge *e*.
- forward(*e*) returns the index of the first edge of the target node of the edge *e*.
- backward(*u*) returns the index of the first incoming edge of node *u*.
- outdegree(*u*) returns the number of outgoing edges outgoing from *u*.
- indegree(*u*) returns the number of incoming edges to *u*.

## 4 Dynamic Functions

Next, we explain the dynamic functions of DynamicBOSS by detail. As previously mentioned, we support edge additions and removals, and node additions and removals. For each of these operations, we need to update the bit vector L, the array of flagged edge labels W, and the position array F.

### 4.1 Utility Procedures

In this section, utility procedures are presented; these procedures are used by the main dynamic operations add-edge and delete-edge.

#### 4.1.1 The add-edge-to-node Procedure

The procedure add-edge-to-node adds an outgoing edge *e* with a specified character *c* to an extant node *u* in the graph. As input, it takes the index pos of the first outgoing edge of *u*, the character *c*, a boolean value indicating if the target node *v* of the edge is already present in *G*, and the index posIn of the first incoming edge to *v* if *v* is present. The procedure consists of two parts; the first handles the addition of the outgoing edge to node *u*, and the second ensures that the target node *v* exists and that the minus flags are correctly set.

We begin by the first part of add-edge-to-node that adds the edge to node *u*. Let *y* be the node label of *u*, that is, the last character of the label of *u*, which is determined by F[edge(*u*)]. If W[pos] = $, then W[pos] is assigned the value *c* and L, F are not modified. Otherwise, the position at which *c* should be inserted is found by incrementing pos and checking the values of W, L. When the position is found, pos is updated to store the position at which to insert. Next, *c* is inserted to W at pos, and 1 is inserted to L at pos or pos + 1, the latter case occurring if and only if pos is the first edge of node *u*. After these insertions, the character *y* is inserted into F at pos. Altogether, these operations add the edge to node *u*. If *c* = $, the procedure terminates; otherwise, execution proceeds to the second part.

The second part of add-edge-to-node considers the target node *v* and ensures minus characters are set correctly, meaning that all incoming edges of node *v* except the first one (in colex order) should have a minus in W. If the node *v* does not exist, its only incoming edge will be *e* and hence the character *c* and not *c*-is correct for *e*. However, if node *v* does exist, then posIn is compared to pos. If pos *≤* posIn, then W[posIn + 1] is set to *c*-. Here, posIn is incremented by one since an edge has been added preceding it. Otherwise, if pos *>* posIn, then W[pos] is set to *c*-.

If *v* needs to be added to the graph *G*, then the index nodeTarget of where the node should be is computed by nodeTarget = L.rank_0_(*F*[*i*_*c*_])+W.rank_*c*_(pos+1), where *i*_*c*_ is the index of the first edge with the last character of its node label equal to *c*. This node index is then converted to an edge index by posTarget = L.select_0_(nodeTarget − 1). Finally, the character $, bit 0, and character *c* are inserted at position posTarget to W, L, and F, respectively. Altogether, these insertions add node *v*, the target of edge *e*, with a dummy outgoing edge to the graph. See Figure A.1 for the illustration of this procedure.

#### 4.1.2 The add-dummy-chain procedure

Procedure add-dummy-chain adds a sequence of dummy edges to the graph. As input, it takes a node label *S*[1, *k* − 1] and a boolean value indicating whether the node with label *S* already exists in the graph.

At the end of the procedure, all *k* − 1 preceding dummy edges to *S*, corresponding to labels

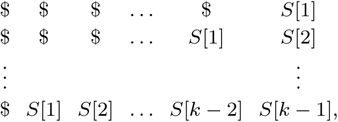

will be present in *G*, along with the required dummy nodes. First, it is necessary to check if *G* contains the dummy node $$ … $, which may be accomplished by checking if F[0] = $. If not, the edge $$ … $ is added to *G* by prepending a $ character to both W and F and prepending a 0 to L. At this point, the dummy node $$ … $ exists, with its first edge at index 0.

Next, the process iteratively, for each edge in the above list, checks if the edge exists by consulting up to four entries of W and L from the current position, and if the edge does exist, follows the edge to the first edge of the next node using the forward operation. Once a node is encountered for which the next outgoing edge does not exist, the add-edge-to-node procedure is called to add the edge to the current node. Once the edge has been added, the forward operation is used to traverse to the next node.

Finally, if the input parameter indicates that the node labeled by *S* already exists in the graph, then prior to the addition of the last edge in the list, the index of the first incoming edge to *S* is determined b y the sequence backward(index-node(*S*)). This information is then passed to add-edge-to-node for the final call to add the last edge in the list above. See Figure A.1 for the illustration of this procedure.

#### 4.1.3 The delete-edge-from-node Procedure

The procedure delete-edge-from-node deletes an outgoing edge *e* from an extant node *u* in the graph. As input, it takes the index pos of the edge within *u* to be deleted, and the label *S* of the edge. The procedure is composed of three parts; the first ensures that the target of the edge has an incoming edge other than *e*, the second ensures that minus flags will be correctly set upon deletion, and the third deletes the outgoing edge from node *u*. Node *u* itself, however, is preserved; if *e* was the only outgoing edge of *u*, it is replaced with a dummy outgoing edge.

First, the target node *v* of the edge is reached by traversing to posTarget = forward(*e*) and then indegree and outdegree are checked. If indegree(*v*) is equal to one and outdegree(*v*) is greater than zero, then add-dummy-chain(*S*[2, *k*]) is called to ensure that *v* has an incoming edge; after this call, *v* will temporarily have two incoming edges until *e* is deleted; also, pos of *e* is recomputed with index after this call. If indegree(*v*) = 1 and outdegree(*v*) = 0, the node *v* is deleted from the graph; this is accomplished directly by removing the character W[posTarget], the bit L[posTarget], and the character F[posTarget]

Next, we ensure that after removal of edge *e* at pos, the minus flags are correctly set. If W[*pos*] = *c*-∈ Σ-, no updates are necessary. Otherwise, we check W to determine if the next occurrence after pos in W of *c* or *c*-is an occurrence of *c*-. If so, this occurrence is replaced with character *c*. The positions of these occurrences are computed using rank, select on W. Finally, the edge is deleted by removing the character W[pos], the bit L[pos], ensuring the node flags for *u* are correct, and removing the character F[pos]. If *e* is the only outgoing edge of *u*, a dummy edge is added to *u* and no change is made to F.

#### 4.1.4 The delete-dummy-chain Procedure

The delete-dummy-chain procedure removes dummy edges and nodes that are no longer needed in the graph. It takes as input the index pos of the first dummy edge to remove and the label *S* of this edge.

The procedure works by calling delete-edge-from-node(pos, *S*) which removes the edge with label *S* from the graph and deletes the target node of this edge if it is isolated. Then, if outdegree(node(pos)) is greater than zero, the procedure terminates. Otherwise, pos is reassigned to backward(node(pos)), the next edge label is computed by $*S*[1, *k* − 1], and the process repeats. This continues until a dummy node with outdegree greater than 0 after removal is encountered.We remark that a dummy node may have only one incoming edge. Altogether this removes all dummy nodes and edges that were only in the graph to support the target of the input edge. Finally, the procedure checks if *W* [0] = 0 and *F* [0] = $; that is, if the edge with label $$ … $ is present in the graph. If it is, it means the dummy node with label $$ … $ has no outgoing edges. In this case, we remove it directly from the graph by removal from W, L, and F at 0.

### 4.2 Edge Addition

An edge *k*-mer *e* with label *S* is added to the graph by procedure add-edge which takes as input the label *S* of *e*. Pseudocode for add-edge is shown in Alg. 1 in the Supplement. The add-edge procedure works as follows: the existence and index of the source node *u*_*e*_ of the edge with label *S*[1, *k*−1] is determined by a call to the operation index-node. If index-node reports that *u*_*e*_ is absent from the graph, then *u*_*e*_ is inserted, along with any necessary dummy nodes and edges, by procedure add-dummy-chain, which is described in §4.1.2. Therefore, by line 5 in the pseudocode, the node *u*_*e*_ is present in the graph and its index is stored in posSource.

Let node *u*_*e*_ correspond to range of edge indices [*f*_*u*_, *l*_*u*_]. Then, by considering W[*f*_*u*_], …, W[*l*_*u*_], it is determined whether *e* already exists in the graph; that is, whether any of these entries equals *S*[*k*] or *S*[*k*]-. If the edge is absent, then procedure add-edge-to-node is called, which updates W, L, and F to add the character *S*[*k*] or *S*[*k*]-to W. This update adds the edge *e* to the graph. The procedure add-edge-to-node is described in detail in §4.1.1. Finally, add-edge-to-node requires knowledge of whether the target node *v*_*e*_ exists, and if so, the index of the first incoming edge to *v*_*e*_. This knowledge is supplied to add-edge-to-node by calls to index-node and backward.

### 4.3 Edge Deletion

An edge *k*-mer *e* with label *S* is deleted from the graph by procedure delete-edge which takes as input the label *S* of *e*. Pseudocode for delete-edge is shown in Alg. 2 in the Supplement. The delete-edge procedure works as follows. First, existence of the edge is checked by a call to index; if it exists, its position is stored in pos. Next, the outdegree is computed of the source node *u* of the edge. This information will be useful to decide if delete-dummy-chain must be called for incoming edges to *u*. Next, the edge *e* is deleted by a call to delete-edge-from-node, which removes the target node as well if it is incident with no other edges. Also, delete-edge-from-node adds dummies to the target if it has outgoing edges but no other incoming edges through a call to add-dummy-chain. Finally, if the edge *e* was the only outgoing edge of *u*, we check for the existence of an incoming dummy edge to *u* by a call to index. If such a dummy edge does exist, we remove this edge, along with *u* itself and any other dummy nodes and edges that are in the graph solely to support *u*, by a call to delete-dummy-chain.

### 4.4 Node Changes

We revisit the definition of the de Bruijn graph and note that the nodes are created and defined as the prefixes and suffixes of the edges. Hence, we do not support the addition and deletion of isolated nodes in the graph and instead, only add or delete a node in the process of addition or deletion of its incident edges. More specifically, addition of a node *u* is accomplished by a call to add-edge for any edge incident with *u*, and deletion of a node *u* is accomplished by deleting every incident edge with *u* through calls to delete-edge as needed.

### 4.5 Time Complexity

We observe that each procedure called by add-edge or delete-edge requires at most *O*(*k*) calls to rank, select, access, insert, and remove on the underlying data structures encoding W, L, and F. Since F is compressed as an array of length 5, it can be accessed and modified in constant time. Since in our implementation we used the library of Prezza (2017) and the alphabet size is constant, the time complexities of each of the five above operations is bounded by *O*(log |*E*|), where |*E*| is the number of edges. Thus, the overall time complexities of add-edge and delete-edge are bounded by *O* (*k* log |*E*|). See Table A.3 in the Supplement for an overview of the running time of these operations.

## 5 Results and Discussion

In this section, we compare the performance of DynamicBOSS to (static) BOSS (Bowe et al., 2012) and the following dynamic de Bruijn graph implementations: FDBG (Crawford et al., 2018), BFT (Holley et al., 2016), and Bifrost (Holley and Melsted, 2019). Hence, we restrict interest to dynamic implementations but note that there exists several space-efficient (colored) de Bruijn graph implementations e.g., (Muggli et al., 2017; Almodaresi et al., 2017; Pandey et al., 2018) and methods that compress the color matrix (Mustafa et al., 2017, 2019; Karasikov et al., 2019). We compared the time required to construct the graph, the size of the de Bruijn graph in memory and stored on hard disk, and the time required to add or delete a *k*-mer as well as the time required to query the *k*-mers. We evaluated these measures on datasets of increasing size. The details of the datasets are described below. We note all experiments were performed on an Intel(R) Xeon(R) CPU E5-2650 v2 2.60 GHz server with 1 TB of RAM, and both resident set size and user process time were reported by the operating system.

### 5.1 Datasets

First, we evaluated the performance using a number of samples from the shotgun metagenomics dataset generated by Noyes et al. (2016) (NCBI SRA accession PRJNA292471). In particular, we used samples 1 and 92 that have 55,242,004 and 11,136,890 reads, respectively, which we refer to as 55M and 11M for simplicity. In addition, we generated smaller samples by randomly selecting 20,000, 200,000 and 2,000,000 sequence reads from sample 92. We refer to these as 20K, 200K and 2M, respectively. Next, we generated larger samples by concatenating samples until the desired length was obtained. Namely, we concatenated samples 1, 2, and 3, which consist of 55,242,004, 44,035,852, and 52,833,978 reads, respectively. We refer to this as sample 150M. We then added samples 4, 5, 9, 22, 25, 33, 34 and 51 to sample 150M to generate the sample 600M.

Second, we evaluated the performance using samples of *E*.*coli*. We used 28,428,648 paired-end reads generated from whole genome sequencing of *E*.*coli* K-12 substr. MG1655 dataset (NCBI SRA accession ERX002508). We refer to this as 28M-e. Next, we split this sample to generate smaller datasets of sizes 20,000, 200,000, 2,000,000, and 14,000,000 reads, which we denote by 20K-e, 200K-e, 2M-e and 14M-e, respectively.

### 5.2 Construction Time and Graph Sizes

We constructed the de Bruijn graph with *k* = 31 for all methods except BFT, where we used *k* = 27 since BFT only accepts values that are multiples of 9. Tables 1 and 2 demonstrate how the construction time and final size of the data structures loaded to memory and stored on disk after construction vary between DynamicBOSS and the competing methods on metagenome and *E*.*coli* datasets, respectively. We did not filter any *k*-mers so that all methods are compared with the same number of *k*-mers. We note that FDBG was not able to construct the graph on datasets larger than 11M on metagenome samples. The reason is that as a hash-based method, it requires a prime value that is large enough with respect to the size of the data, hence when the data gets very large, and assigning the respective large prime is not trivial, this method fails due to the possibility of collisions. NA is given in Table 1 for the entries that FDBG failed.

**Table 1.** Comparison of construction time (hh:mm:ss), size of the graphs in the memory (MB or GB) and disk (Bits/k-mer). The graphs are built on metagenome samples of different sizes. 20K, 200K, 2M, 11M, 55M, 150M, and 600M are samples with 20,000, 200,000, 2,000,000, 11,136,890, 55,242,004, 152,111,834 and 589,340,896 reads and 1,353,272, 13,430,422, 126,032,942, 592,287,827, 1,778,630,301, 4,821,975,335 and 15,705,461,763 solid 31-mers respectively. N/A is shown where FDBG could not build the graph. Our method is highlighted in bold.

|  | BOSS |  |  | DynamicBOSS |  |  | BFT |  |  | FDBG |  |  | Bifrost |  |  |
| --- | --- | --- | --- | --- | --- | --- | --- | --- | --- | --- | --- | --- | --- | --- | --- |
|  | Disk | Memory | Time | Disk | Memory | Time | Disk | Memory | Time | Disk | Memory | Time | Disk | Memory | Time |
| 20K | 2.9 | 0.5 MB | 00:00:01 | **7.0** | **1.2 MB** | **00:00:01** | 48.7 | 16.5 MB | 00:00:02 | 17.1 | 7.8 MB | 00:00:05 | 6.3 | 19.0 MB | 00:00:08 |
| 200K | 2.8 | 4.7 MB | 00:00:07 | **7.1** | **12.3 MB** | **00:00:22** | 46.4 | 92.0 MB | 00:00:12 | 17.2 | 41.4 MB | 00:00:59 | 6.4 | 120.2 MB | 00:01:15 |
| 2M | 2.7 | 43.3 MB | 00:01:16 | **6.7** | **112.9 MB** | **00:03:31** | 45.6 | 835.5 MB | 00:02:26 | 16.7 | 348.3 MB | 00:11:47 | 7.3 | 1.8 GB | 00:11:16 |
| 11M | 2.6 | 197.6 MB | 00:06:04 | **6.4** | **514.3 MB** | **00:17:16** | 44.0 | 834.5 MB | 00:11:09 | 16.2 | 1.5 GB | 01:27:28 | 10.0 | 7.6 GB | 00:59:25 |
| 55M | 2.6 | 579.0 MB | 00:18:29 | **6.2** | **1.4 GB** | **00:53:12** | 42.0 | 9.0 GB | 00:39:30 | NA | NA | NA | 13.8 | 16.2 GB | 03:10:59 |
| 150M | 2.6 | 1.5 GB | 01:00:25 | **6.1** | **4.0 GB** | **02:44:50** | 42.6 | 23.3 GB | 02:57:28 | NA | NA | NA | 14.9 | 65.6 GB | 09:04:39 |
| 600M | 2.5 | 5.0 GB | 03:38:29 | **6.1** | **12.7 GB** | **09:21:17** | 39.6 | 67.3 GB | 13:02:42 | NA | NA | NA | 19.8 | 138.2 GB | 40:36:55 |

**Table 2.** Comparison of construction time (hh:mm:ss), size of the graphs in the memory (MB or GB) and disk (Bits/*k*-mer). Graphs are built on E.coli samples of different sizes. 20K, 200K, 2M, 14M and 28M are samples with 20,000, 200,000, 2,000,000, 14,000,000, and 28,428,648 E.coli reads and 1,310,799, 8,395,324, 28,913,955, 150,906,786, 243,947,156 solid *k*-mers respectively. Our method is highlighted in bold.

|  | BOSS |  |  | DynamicBOSS |  |  | BFT |  |  | FDBG |  |  | Bifrost |  |  |
| --- | --- | --- | --- | --- | --- | --- | --- | --- | --- | --- | --- | --- | --- | --- | --- |
|  | Disk | Memory | Time | Disk | Memory | Time | Disk | Memory | Time | Disk | Memory | Time | Disk | Memory | Time |
| 20K-e | 2.9 | 0.4 MB | 00:00:01 | **7.3** | **1.2 MB** | **00:00:01** | 48.7 | 16.9 MB | 00:00:02 | 17.0 | 7.5 MB | 00:00:06 | 6.8 | 19.1 MB | 00:00:10 |
| 200K-e | 2.5 | 2.6 MB | 00:00:05 | **6.0** | **6.8 MB** | **00:00:13** | 47.6 | 49.1 MB | 00:00:05 | 16.1 | 23.7 MB | 00:00:37 | 13.0 | 65.7 MB | 00:00:55 |
| 2M-e | 2.5 | 9.2 MB | 00:00:16 | **5.8** | **21.8 MB** | **00:00:40** | 45.8 | 151.7 MB | 00:00:25 | 15.4 | 67.7 MB | 00:02:26 | 29.8 | 328.8 MB | 00:04:45 |
| 14M-e | 2.7 | 50.7 MB | 00:01:31 | **5.7** | **115.2 MB** | **00:04:13** | 43.4 | 867.2 MB | 00:02:48 | 15.3 | 309.3 MB | 00:26:46 | 36.8 | 2.4 GB | 00:40:31 |
| 28M-e | 2.7 | 82.8 MB | 00:03:00 | **5.7** | **187.0 MB** | **00:07:07** | 43.4 | 1.3 GB | 00:05:12 | 15.3 | 498.2 MB | 00:30:54 | 38.4 | 2.6 GB | 01:20:23 |

As shown in these tables, BOSS required the least amount of space when stored on disk and had the most efficient construction. It required less than 3 hours and 40 minutes to construct the graph on over 600M reads. And each graph required less than 2.9 bits / *k*-mer. With the addition of dynamic features, DynamicBOSS requires between 5.7 and 7.0 bits / *k*-mer, giving an increase of approximately 5 bits / *k*-mer. Bifrost required between 6.3 and 38.4 bits / *k*-mer. Interestingly, its size increased as the sample size increased, e.g., the space is 6.3 bits / *k*-mer for the 20K sample and 19.8 bits / *k*-mer for the 600M sample. Bifrost was smaller on only the two smallest metagenome samples (20K and 200K) and on the smallest *E*.*coli* sample (20K-e). BFT and FDBG had between 39.6 and 48.7, and 16.2 and 17.2 bits / *k*-mer. Therefore, DynamicBOSS required the least amount of space to store the graph on samples with more than 200K reads, and this space consistently decreased as the sample size increased.

In addition, Bifrost required between 3 and 11 times more time than DynamicBOSS to construct the graph. For example, on 600M and 28M-e samples, Bifrost required 40 hours 36 minutes and 1 hour 20 minutes, and DynamicBOSS required less than 9.5 hours and 7 minutes. BFT had comparable runtime with DynamicBOSS – it was slightly more efficient for smaller samples (less than 55M reads) and DynamicBOSS was slightly more efficient for the larger samples. FDBG was comparable to Bifrost but as mentioned above, it was unable to construct the graphs on metagenome samples with more than 11M reads.

Lastly, Memory in Tables 1 and 2 gives the size of the graphs when loaded to RAM. As shown, DynamicBOSS required the least amount of memory among all dynamic methods. This measure is important for dynamic structures since updating the graph in all of the tools is memory resident. FDBG required the second-least amount of memory but could not be constructed on larger samples. The sizes of BFT and Bifrost graphs in memory were up to 7 times and 16 times (respectively) larger than that of DynamicBOSS. For instance, while the size of DynamicBOSS on 600M and 28M-e was 12.7 GB and 187.0 MB, BFT used 67.3 GB and 1.3 GB, and Bifrost used 138.2 GB and 2.6 GB respectively.

### 5.3 Deletion and Addition of *k*-mers

Next, we compared the time required by DynamicBOSS, Bifrost and FDBG to add random *k*-mers to the de Bruijn graph. In theory, BFT is capable of this but it is no longer supported or functional (Holley, 2019). For this comparison, we randomly selected 4,847,680 *k*-mers from two strains of salmonella (ERR044710 and ERR044711 from NCBI BioProject PRJNA18384) and added them to the de Bruijn graphs constructed in the previous section. For example, these *k*-mers were added to the graphs constructed on 20K sample, 200K sample, 2M, and so forth. We then calculated the mean time of the 4,847,680 additions across the 7 metagenome samples and the 6 *E*.*coli* samples. Figures 2a and 2b illustrate these means. We note that the values on the mantissa of these plots show the size of the original graph (before performing addition), e.g 20K to 600M on metagenome samples, and 20K to 28M on *E*.*coli*, and number of added *k*-mers to graphs in this experiment is constant and equal to 4,847,680. Although FDBG and Bifrost have more efficient addition time, the addition time of Bifrost increased dramatically as the size of the original graph (before addition) grows. For instance while the average time per *k*-mer addition on 2M graph was 1.38e-05 (s), this value increased to 9.48e-04 (s) on 600M graph. Also, FDBG could only be evaluated on smaller datasets since the graph could not be constructed for large samples (e.g *>* 11M metagenome reads)–see Section 5.2 for the discussion. Hence, the addition time of FDBG on large graphs is unknown.

**Fig. 2:**
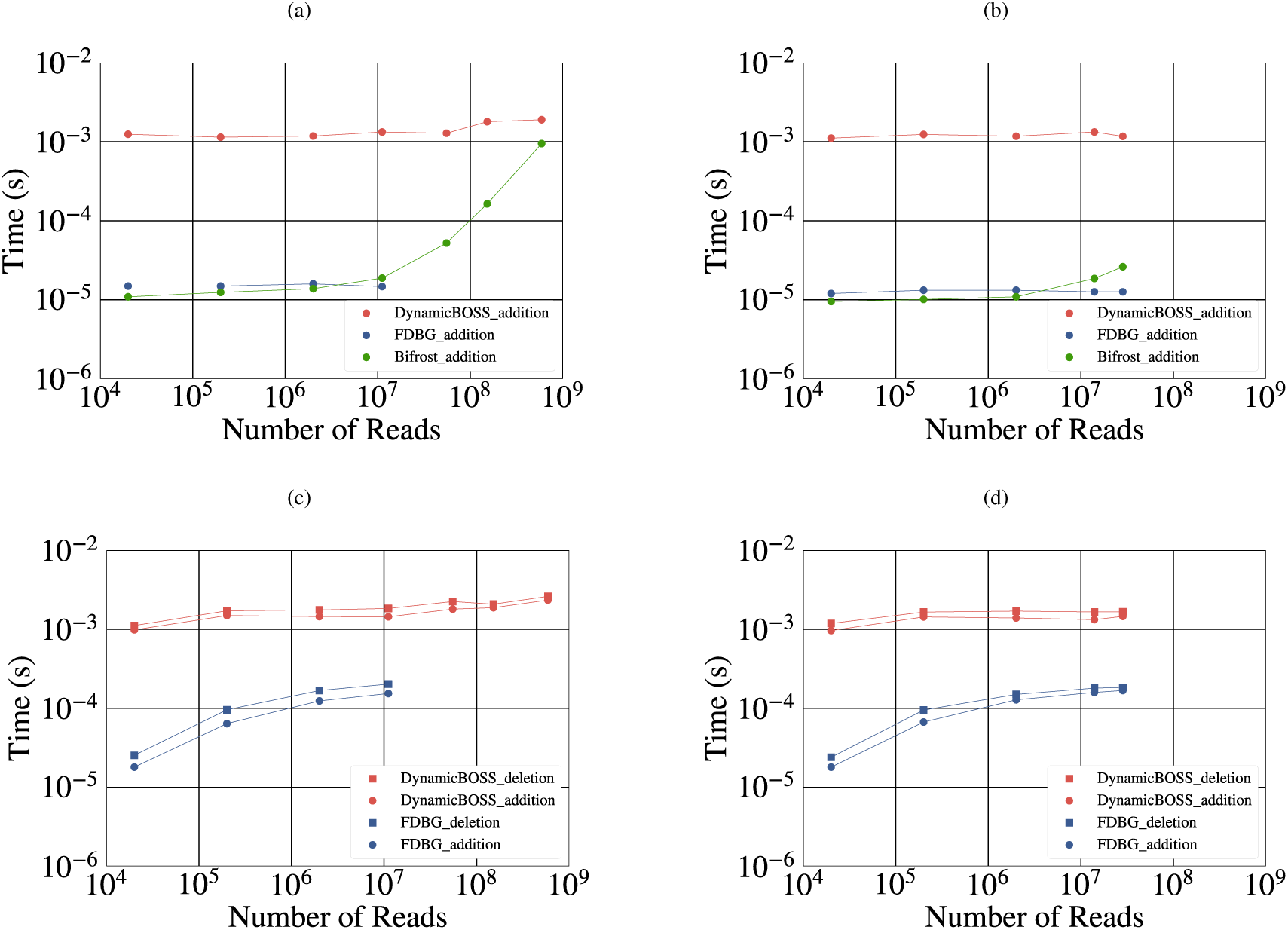
An Illustration of the average required time to add random *k*-mers to the de Bruijn graphs built on (a) metagenome and (b) *E. coli* samples, and the time required to delete and re-add *k*-mers from the de Bruijn graphs built on (c) metagenome and (d) *E. coli* samples. We have used the discussed graphs in Section 5.2 in these experiments. We note that FDBG could not add or build on samples with ⩾ 55M reads and therefore, the corresponding data points are missing from (a) and (c).

Lastly, we evaluated the time to delete and re-add *k*-mers from/to the de Bruijn graph. Here, we compared our method to FDBG since it is the only other method that supports deletion. Again, we deleted/added to the de Bruijn graphs constructed in the previous section using the metagenome and *E. coli* datasets. For each graph, we randomly selected 1,000,000 *k*-mers from each of these de Bruijn graphs, first deleted and then re-added them back to the graph. Figures 2c and 2d show the mean deletion and addition time on the metagenomic and *E. coli* datasets, respectively. Again, FDBG had faster deletion time but it could only support building the small graphs with *<* 55M reads. Hence, its dynamic operation time on large graphs is unknown.

In summary, Bifrost has more efficient addition time than our method but cannot support deletion and the addition time increased significantly as the size of the graph increased, FDBG had more efficient addition and deletion time but is only applicable to small samples.

### 5.4 Querying *k*-mers

Finally, we compared the time required by DynamicBOSS, BOSS, Bifrost, BFT and FDBG to query *k*-mers. First, we selected 1,000,000 *k*-mers from the set of available edges in the de Bruijn graphs constructed in Section 5.2 uniformally at random without replacement and measure the average time for querying them in the same graph. Next, we randomly selected 4,847,680 *k*-mers from two strains of salmonella (ERR044710 and ERR044711 from NCBI BioProject PRJNA18384), and measure the average time for querying them in the de Bruijn graphs. Figure 3a and 3b show these results in graphs built on metagenome and *E*.*coli* samples respectively. We note that the values on the mantissa of these plots show the size of the original graph, e.g 20K to 600M on metagenome samples, and 20K to 28M on *E*.*coli*, and numbers of queried *k*-mers in graphs in these experiments are constant and equal to 1,000,000 member *k*-mers and 4,847,680 random *k*-mers.

**Fig. 3:**
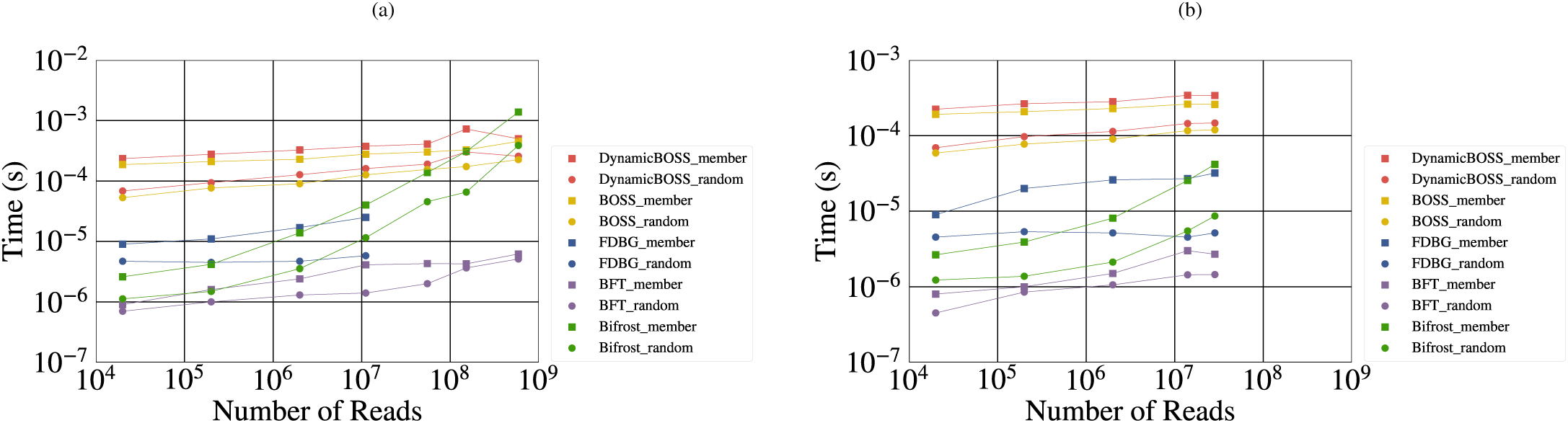
An Illustration of the average required time to query 1,000,000 random *k*-mers that were known to be contained in the de Bruijn graph, and to query 4,847,680 random *k*-mers taken from the Salmonella genome. We show the results of the former experiment with extension *member* and the latter experiment with extension *random* in these plots. These experiments were performed in de Bruijn graphs constructed on (a) metagenome samples and E.coli samples mentioned in Section 5.2. We note that FDBG could not build or query on samples with ⩾ 55M reads and therefore, the corresponding data points are missing from (a).

In summary, BFT has the most efficient querying time among all tools. The next most efficient querying belongs to FDBG, which is unable to construct and query on samples ⩾ 55M reads–see Section 5.2 for the discussion. Hence, Figure 3a does not include the corresponding data points. The query time of DynamicBOSS is slightly more than BOSS, and finally even though Bifrost has efficient querying time for smaller samples, its efficiency drops as the sample sizes grow and it has the least efficient query time for the largest sample which is 600M.

## 6 Conclusion

A dynamic de Bruijn graph is attractive for both de Bruijn graphs and colored de Bruijn graphs. It is standard practise in assemblers to remove structures such as tips and bubbles caused by sequencing errors from the graph. If the graph representation is static then it has to be rebuilt or a separate data structure has to be used to track deleted edges and nodes. Therefore, a dynamic de Bruijn graph would significantly streamline this cleaning step. Being able to dynamically update the colored de Bruijn graph is also extremely attractive because it enables a stored graph to be updated as a set of sequence reads is added or removed from a dataset without reconstruction, and allows analysis of a selected number of samples. Yet, it is challenging to design data structures that are both space-efficient and mutable. Here, we showed that we can take advantage of recent advancements in dynamic data structures and build the de Bruijn graph in a manner that is space-efficient and mutable. Our experimental results showed that with a small increase in the space usage (approximately 5 bits / *k*-mer), we can achieve a fully dynamic data structure for the de Bruijn graph.

## A.1 Supplement

### A.1.1 Comparison of BOSS and DynamicBOSS

Table A.3 shows the time complexity of different functions in Non-dynamic BOSS and DynamicBOSS, where *k* is the *k*-mer size and *E* is the number of the edges in the graph.

**Table A.3.**
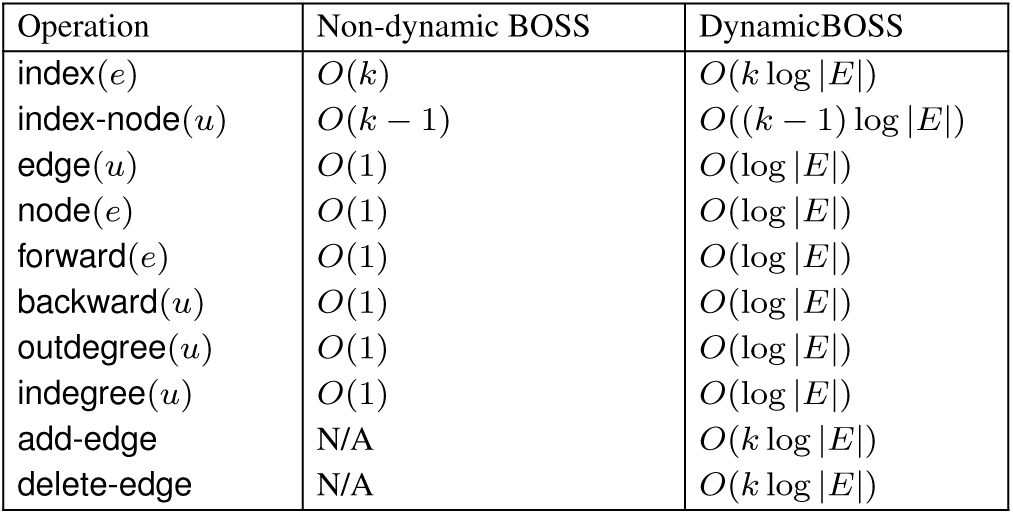
Comparison of the running time of operations of non-dynamic BWT-based representation of the de Bruijn graph (BOSS) and the dynamic version presented in this paper. *E* denotes the number of edges and *k* denotes the *k*-mer value.

### A.1.2 Pseudocode for Dynamic Functions

#### Algorithm 1 Add edge with label *S* to graph

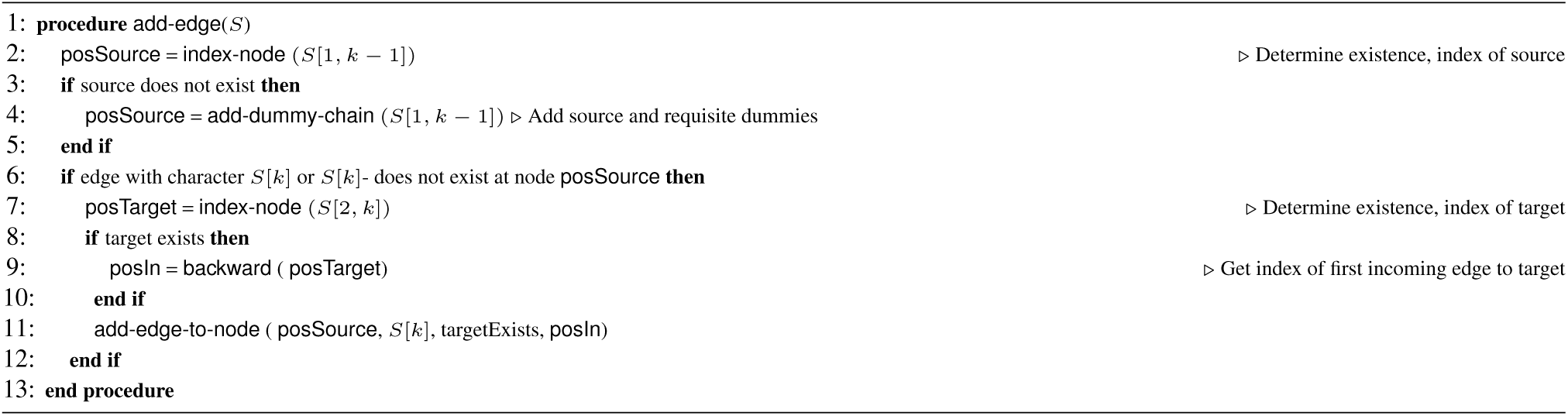

#### Algorithm 2 Delete edge with label *S* from graph

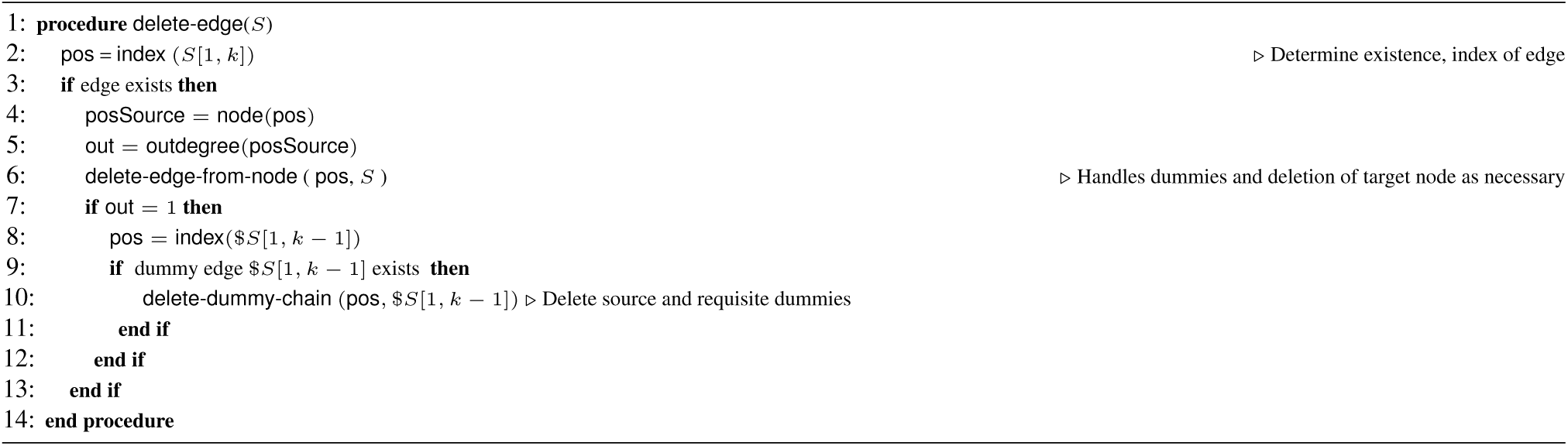

### A.1.3 An illustration of add-edge procedure

We add an edge *e* with label *S* = `ATGA` to the graph *G* of Figure 1 in Section 3 and show all steps of the algorithm. Note that here *u*_*e*_ = `ATG` and *v*_*e*_ = `TGA` are the source node and the sink node of edge *e* respectively. First using the operation index-node (*u*_*e*_) the existence of the source node is checked. Since *u*_*e*_ is absent, it must be inserted to *G* along with any necessary dummy nodes and edges by procedure add-dummy-chain (*u*_*e*_). This procedure starts with checking if the node $$$ is already in the graph. Since F[0] = $ or F[$] = 0 in our compressed representation of F, this node is present in the graph and its posSource is 0. Hence iteratively for each edge $$$`A`, $$`AT` and $`ATG` it checks if the edge exists and if not, it adds the edge to the corresponding source node by calling the procedure add-edge-to-node. We show the whole process step by step in Figure A.1 highlighting the changes by the first and second step of procedure add-edge-to-node in red and grey respectively. Also, the updates on F are colored in blue. In Figure A.1(1), we first check if $$$`A` exists or not. The range of source node $$$ is [0,0]. Note that this range is found as follows: [L.select_0_[0], L.select_0_[1] − 1]. W[0] ≠ `A` or `A`− shows that the edge $$$`A` is absent and should be added to the graph. This is accomplished by calling add-edge-to-node (0, `A`, 0, null). Note that 0 as the third input shows that the target node of this edge which is $$`A` is not present in the graph. The first step of add-edge-to-node adds the edge $$$`A` to the node $$$. In order to do this, with comparing the edge label `A` with the edge label at posSource which is W[0] = `C`, the position to insert the edge is finalized which in this case is position 0. Next, `A` is inserted at position 0 in W. L[0] remains unchanged since now $$$`A` is the first outgoing edge of $$$, and a 1 is inserted at L[1] as the edge $$$`C` is no longer the first outgoing edge of $$$. Finally, the first occurrence of every character after $ is incremented by one in F. Next the second part of add-edge-to-node is executed. Since the target node $$`A` is absent, we find the nodeTarget as follows: nodeTarget = L.rank_0_(F[*i*_*c*_]) + W.rank_*c*_(pos + 1). With replacing F[*i*_*c*_] = 2 and pos = 0, nodeTarget = 1+1 = 2 and posTarget = L.select_0_(nodeTarget − 1) = 2 are the node index and the corresponding edge index of the target node. Finally the character $ and bit 0 are added to W and L at position 2 and the first occurrence of every character after `A` will get incremented by one in F – note that this step is required since every node should have at least one outgoing edge. Next we need to add the edge $$`AT` to the graph. In Figure A.1(2) add-edge-to-node (2, `T`, 0, null) is called and the edge label `T` and bit 0 are added at position 2 of W and L respectively, and F remains unchanged. Due to the absence of the target node $`AT`, it should be added to the graph as well. This time nodeTarget = 12 + 1 = 13 and posTarget = 13. Hence the character $ and bit 0 are added to W and L at position 13, and since the first position of occurrence of `T` is still at position 13, F remains unchanged. In Figure A.1(3), add-edge-to-node (13, `G`, 0, null) is called to add the edge $`ATG` to the graph. Hence the character `G` and bit 0 are added to W and L at position 13 and F is unchanged. This time nodeTarget = 8 + 4 = 12 and posTarget = 12. Hence the character $ and bit 0 are added to W and L at position 12. Note that since posTarget *<* posSource, the updates of the first step (colored in red) are shifted to position 14. Finally the first occurrence of characters after `G` (only `T`) is incremented by one in F. In Figure A.1(4) add-edge-to-node (12, `A`, 1, 13) is called. Note that the target node which is `TGA` is present in the graph and is located at position 13. Now after adding the character `A` and bit 0 to W and L at position 12 and keeping F unchanged, the second step of add-edge-to-node is executed which this time takes care of the minuses. Since posTarget *>* posSource, we add a minus to the edge label of posTarget, since it is not the first incoming edge to the node `TGA` anymore.

**Fig. A.1:**
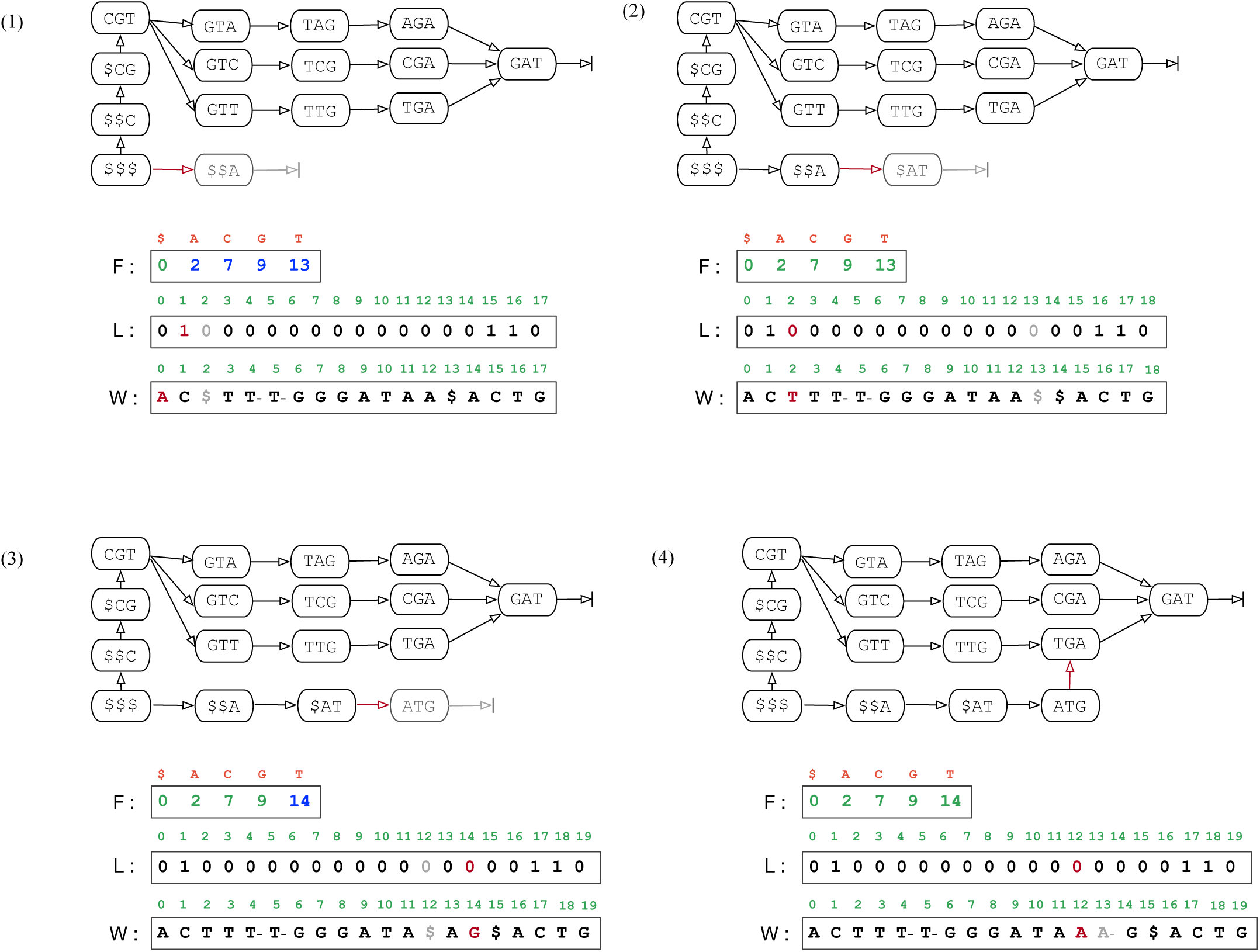
The addition of edge *e* with label *S* = `ATGA` to the graph *G* of Figure 1 which is built on sequences CGTAGAT, CGTCGAT, CGTTGAT. Note that BOSS stores only three vectors F, L and W. Addition of edges $$$`A`, $$`AT` and $`ATG` and `ATGA` is shown in (1), (2), (3), and (4) respectively. The changes made by the first and second step of procedure add-edge-to-node is highlighted in colors red and grey respectively.

## References

Almodaresi, F., Pandey, P., and Patro, R. (2017). Rainbowfish: A succinct colored de Bruijn graph representation. In: Leibniz International Proceedings in Informatics (LIPIcs), 88:18: 1–18: 15.

Bowe, A., Onodera, T., Sadakane, K., and Shibuya, T. (2012). Succinct de Bruijn graphs. In International Workshop on Algorithms in Bioinformatics (WABI), pages 225–235. Springer.

Burrows, M. and Wheeler, D. (1994). A block sorting lossless data compression algorithm. Technical Report 124, Digital Equipment Corporation, Palo Alto, California.

Chikhi, R. and Rizk, G. (2013). Space-efficient and exact de Bruijn graph representation based on a Bloom filter. Algorithms for Molecular Biology., 8(22).

Conway, T. C. and Bromage, A. J. (2011). Succinct data structures for assembling large genomes. Bioinformatics, 27(4):479–486.

Cordova, J. and Navarro, G. (2016). Practical dynamic entropy-compressed bitvectors with applications. In International Symposium on Experimental Algorithms, pages 105–117.

Crawford, V., Kuhnle, A., Boucher, C., Chikhi, R., and Gagie, T. (2018). Practical dynamic de bruijn graphs. Bioinformatics, 34(24):4189–4195.

Ferragina, P. and Manzini, G. (2005). Indexing compressed text. J. ACM, 52(4):552–581.

Grossi, R. et al. (2013). Dynamic compressed strings with random access. In International Colloquium on Automata, Languages, and Programming, pages 504–515.

Holley, G. (2019). Personal email communication with authors of BFT.

Holley, G. and Melsted, P. (2019). Bifrost–highly parallel construction and indexing of colored and compacted de bruijn graphs. bioRxiv.

Holley, G., Wittler, R., and Stoye, J. (2016). Bloom filter trie: an alignment-free and reference-free data structure for pan-genome storage. Algorithms for Molecular Biology, 11.

Iqbal, Z., Caccamo, M., Turner, I., Flicek, P., and McVean, G. (2012). De novo assembly and genotyping of variants using colored de Bruijn graphs. Nature Genetics, 44(2):226–232.

Karasikov, M., Mustafa, H., Joudaki, A., Javadzadeh No, S., Rätsch, G., and Kahles, A. (2019). Sparse binary relation representations for genome graph annotation. In: Cowen L. (eds) Research in Computational Molecular Biology. RECOMB 2019. Lecture Notes in Computer Science, 11467:120–135.

Klitzke, P. and Nicholson, P. (2016). A general framework for dynamic succinct and compressed data structures. Proceedings of the 18th ALENEX, pages 160–173.

Manber, U. and Myers, G. W. (1993). Suffix arrays: A new method for on-line string searches. SIAM Journal on Computing, 22(5):935–948.

Muggli, M., Alipanahi, B., and Boucher, C. (2019). Building large updatable colored de bruijn graphs via merging. Bioinformatics, 35(14):i51–i60.

Muggli, M., Bowe, A., Noyes, N., Morley, P., Belk, K., Raymond, R., Gagie, T., Puglisi, S., and Boucher, C. (2017). Succinct colored de bruijn graphs. Bioinformatics, 33(20):3181–3187.

Mustafa, H., Kahles, A., Karasikov, M., and Rätsch, G. (2017). Metannot: A succinct data structure for compression of colors in dynamic de Bruijn graphs. BioRxiv.

Mustafa, H., Schilken, I., Karasikov, M., Eickhoff, C., Rätsch, G., and Kahles, A. (2019). Dynamic compression schemes for graph coloring. Bioinformatics, 35(3):407–414.

Mäkinen, V. and Navarro, G. (2006). Dynamic entropy-compressed sequences and full-text indexes. In: Lewenstein M., Valiente G. (eds) Combinatorial Pattern Matching (CPM), 4009.

Navarro, G. and Nekrich, Y. (2014). Optimal dynamic sequence representations. SIAM Journal on Computing, 43(5):1781–1806.

Noyes, N. et al. (2016). Resistome diversity in cattle and the environment decreases during beef production. eLife, 5:e13195.

Pandey, P., Almodaresi, F., Bender, M., Ferdman, M., Johnson, R., and Patro, R. (2018). Mantis: A fast, small, and exact large-scale sequencesearch index. Cell Systems, 7(2):201–207.

Pevzner, P., Tang, H., and Waterman, M. (2001). An eulerian path approach to DNA fragment assembly. Proceedings of the National Academy of Sciences (PNAS), 98(17):9748–9753.

Prezza, N. (2017). A framework of dynamic data structures for string processing. In International Symposium on Experimental Algorithms (SEA), pages 11:1–11:15. Leibniz International Proceedings in Informatics (LIPIcs).

Simpson, J. et al. (2009). ABySS: A parallel assembler for short read sequence data. Genome Research, 19(6):1117–1123.

Álvarez García, S. et al. (2019). Compact and efficient representation of general graph databases. Knowledge and Information Systems, 60(3):1479–1510.

